# Volatile Metabolic Signatures Reveal Early Pulmonary Responses to PFAS Exposure

**DOI:** 10.64898/2026.09.23.753921

**Authors:** Patrica-Ivy Agorsor, Ta-Chun Lin, Yue-Wern Huang, Michael O. Eze

## Abstract

Inhalation of airborne per- and polyfluoroalkyl substances (PFAS) represents an important yet understudied exposure route. While volatile organic compounds (VOCs) have been widely explored as biomarkers of toxicant exposure, their potential for early detection of PFAS-induced pulmonary toxicity remains largely unexplored. This study investigated VOC profiles in human alveolar (A549) and bronchial (BEAS-2B) epithelial cells following exposure to environmentally relevant PFAS concentrations (1, 5, and 10 ppb) for 12, 24, and 48 h. VOCs were analyzed using headspace solid-phase microextraction coupled with gas chromatography-Orbitrap mass spectrometry. We observed significant concentration- and time-dependent alterations in VOC profiles in both cell models. The principal VOCs associated with PFAS exposure include butanal, acetone, 2-butanone, 2-nonanone, (E)-2-nonenal, (E)-2-octenal, (E)-2-heptenal, 2,4-heptadienal, 2,4-nonadienal, 2,4-decadienal, 1-pentanol, and 1-hexanol. Cell type-specific regulations were observed, reflecting differential metabolic responses between transformed and non-transformed lung epithelial cells. Metabolite set enrichment analysis indicated perturbations in ketone body metabolism, consistent with altered cellular energy metabolism. Complementary qPCR analysis revealed altered expressions of CYP1A1, CYP2E1, ALDH1A1, and ALDH2, supporting a mechanistic link between PFAS exposure and oxidative stress, lipid peroxidation, and impaired aldehyde detoxification. These findings demonstrate that VOC profiling, combined with transcriptional analysis, can reveal PFAS-induced metabolic perturbations in pulmonary epithelial cells.

## 1. Introduction

Per- and polyfluoroalkyl substances (PFAS), often referred to as “forever chemicals”, are synthetic chemicals characterized by carbon chains that are fully (perfluoro-) or partially (polyfluoro-) fluorinated, often terminated with functional groups such as carboxylic acid, sulfonic acid, or amine.^1^ Their exceptional thermal stability, chemical resistance, and amphiphilic nature have led to their widespread use in industrial processes and everyday consumer products, including firefighting foams, nonstick cookware, stain-resistant fabrics, and food packaging materials.^2^ These properties contribute to the extreme environmental persistence, widespread distribution, and bioaccumulation in humans and wildlife, raising public health concerns.^3^ Human exposure to PFAS has primarily been attributed to contaminated drinking water and food.^4^ However, other exposure pathways, including inhalation of airborne PFAS, may also contribute to human exposure, particularly in occupational and indoor environments, although it remains a less well-characterized exposure route^5^. Volatile PFAS precursors, including fluorotelomer alcohols (FTOHs), perfluoroalkyl sulfonamides, and related compounds, are commonly detected in indoor air and dust.^4,6-8^ Once inhaled, these compounds may be absorbed into the body, contributing to the overall PFAS body burden and associated health risks.^9^ PFAS can interfere with multiple biological pathways, including lipid, glucose, bile acid, and amino acid metabolism, as well as hormonal regulation and gut microbiome composition.^10-12^ In addition, PFAS exposure has been associated with a wide range of adverse health outcomes, including immunotoxicity,^13,14^ liver and kidney dysfunction,^15-17^ endocrine disruption,^18^ reproductive and developmental effects,^19-21^ and an increased risk of certain cancers.^22-24^

Despite growing evidence of PFAS toxicity, current biomonitoring approaches remain largely focused on measuring PFAS concentrations in blood, serum, or plasma using liquid chromatography-mass spectrometry (LC-MS). While these methods provide valuable information regarding internal PFAS burden, they are invasive, costly, and often fail to capture early biochemical and physiological responses to exposure.^12,25^ Efforts to identify secondary biomarkers of PFAS exposure have explored inflammatory markers,^26^ oxidative stress indicators,^27,28^ and metabolic parameters.^29,30^ Although informative, these biomarkers generally lack specificity and may not adequately reflect early exposure-induced metabolic alterations, particularly following inhalation exposure. Therefore, there remains a need for sensitive and non-invasive approaches capable of detecting early biological responses associated with PFAS exposure.

Volatile organic compounds (VOCs) represent a promising class of non-invasive early exposure biomarkers because they reflect real-time and dynamic metabolic and physiological processes.^31,32^ VOC profiling has been successfully applied to monitor exposure to environmental contaminants and to detect disease-associated metabolic changes prior to conventional detection methods.^33-40^ Furthermore, advances in analytical instrumentation have enabled the detection of VOCs at trace concentrations in a variety of biological and environmental matrices.^41^ Because the respiratory system represents the primary interface for airborne PFAS exposure, VOC-based metabolomics may provide valuable insight into PFAS-induced metabolic alterations within the pulmonary tissues. Exposure to PFAS may induce distinct VOC signatures that may reflect underlying biochemical responses. However, to the best of our knowledge, no studies have investigated and characterized the VOC metabolic responses of the human alveolar epithelial cells and the human bronchial epithelial cells following PFAS exposure.

Therefore, this study investigated VOC metabolic responses in A549 and BEAS-2B cells following exposure to a mixture of environmentally relevant PFAS compounds including perfluorobutanoic acid (PFBA), perfluoropentanoic acid (PFPeA), perfluorohexanoic acid (PFHxA), perfluoroheptanoic acid (PFHpA), perfluorooctanoic acid (PFOA), perfluorononanoic acid (PFNA), perfluorodecanoic acid (PFDA), perfluorobutanesulfonic acid (PFBS), perfluorohexanesulfonic acid (PFHxS), and perfluorooctanesulfonic acid (PFOS)^42-45^ at different concentrations (1 ppb, 5 ppb, and 10 ppb). VOC profiles were evaluated at 12, 24, and 48 h post-exposure time points using headspace solid-phase microextraction coupled with Orbitrap gas chromatography-mass spectrometry (HS-SPME-GC-Orbitrap MS). Further, quantitative PCR analysis was performed to evaluate alterations in genes involved in xenobiotic metabolism and detoxification pathways. By integrating volatilomic and transcriptional responses, this study aimed to provide insight into the metabolic response of PFAS exposure in pulmonary epithelial cells and to identify potential VOC biomarkers associated with inhalation exposure.

## 2. Materials and Methods

### 2.1 Chemicals and reagents

The human alveolar epithelial carcinoma cell line A549 (ATCC^®^ CCL-185™) and the human bronchial epithelial cell line BEAS-2B (ATCC^®^ CRL-9609™) were obtained from the American Type Culture Collection (ATCC, Manassas, VA, USA). A549 cells were cultured in Ham’s F-12 medium, whereas BEAS-2B cells were cultured in DMEM/F-12. Fetal calf serum (FCS) and penicillin-streptomycin were purchased from Fisher Scientific (Pittsburgh, PA, USA). Cells were maintained at 37°C in a humidified incubator with 5% CO_2_. A native per- and polyfluoroalkyl substances (PFAS) standard mixture containing perfluorobutanoic acid (PFBA), perfluoropentanoic acid (PFPeA), perfluorohexanoic acid (PFHxA), perfluoroheptanoic acid (PFHpA), perfluorooctanoic acid (PFOA), perfluorononanoic acid (PFNA), perfluorodecanoic acid (PFDA), perfluorobutane sulfonate (PFBS), perfluorohexane sulfonate (PFHxS), and perfluorooctane sulfonate (PFOS) was obtained from Wellington Laboratories Inc. (Guelph, ON, Canada). Sodium chloride (NaCl) was purchased from Sigma-Aldrich (St. Louis, MO, USA). Volatile organic compounds (VOCs) were extracted using a fiber 80 µm divinylbenzene/carboxen/polydimethylsiloxane (DVB/CAR/PDMS) solid-phase microextraction (SPME) fiber purchased from Thermo Fisher Scientific (Waltham, MA, USA).

### 2.2 Cell culture and PFAS exposure

A549 and BEAS-2B cells were cultured in Ham’s F-12 and DMEM/F-12 medium, respectively, and supplemented with 10% fetal calf serum (FCS) and 100 U/mL penicillin-streptomycin. The cells were maintained at 37 °C in a humidified incubator with 5% CO_2_. The cells were seeded at a density of 1 × 10^5^ cells per 60 mm culture dish. After allowing the cells to attach for 16 hours, the culture medium was replaced with medium containing 0, 1, 5, or 10 parts per billion (ppb) of the PFAS mixture. Exposure was conducted in 60 mm culture dishes at 12 h, 24 h, and 48 h time points, with biological replicates (n = 3) for each condition. In parallel, control wells containing cells in complete medium without PFAS (“control cells”) and matrix blank wells containing only complete medium with PFAS (no cells) were prepared to assess background interaction and baseline signal. Following exposure, culture supernatants were collected and immediately stored at −80 °C until analysis.

### 2.3 Metabolite extraction and gas chromatography-mass spectrometry (GC-MS) analysis

A 5 mL aliquot of thawed culture supernatant from each condition was transferred into 20 mL headspace vials, followed by the addition of 2.0 g sodium chloride (NaCl) to enhance the release of volatile analytes through the salting-out effect. Volatile organic compounds were extracted using headspace solid-phase microextraction (HS-SPME) with a TriPlus RSH autosampler (Thermo Fisher Scientific, Waltham, MA, USA). Samples were incubated at 65 °C for 10 min under continuous agitation, followed by extraction for 10 min using a preconditioned 80 μm DVB/CAR/WR/PDMS triphasic SPME fiber. The fiber was thermally desorbed in the GC injector at 225 °C for 3 min. GC-MS analysis was performed using an Orbitrap Exploris GC–MS system (Thermo Fisher Scientific, Waltham, MA, USA) equipped with a TG-5SilMS capillary column (30 m × 0.25 mm i.d. × 0.25 μm film thickness). Ultra-high-purity helium (Airgas, USA) was used as the carrier gas at a constant flow rate of 1.2 mL/min, with a front inlet purge flow of 5.0 mL/min. Samples were analyzed in splitless mode. The GC oven temperature program was as follows: an initial temperature of 40 °C held for 1 min, followed by an increase to 280 °C at a rate of 25 °C/min and held for 6 min. The injector, transfer line, and ion source temperatures were maintained at 280 °C. The mass spectrometer was operated in positive electron ionization (EI) mode at 70 eV, and full-scan mass spectra were acquired over an m/z range of 50–700.

### 2.4 Cytotoxicity Assay

This study employed the 3-(4,5-dimethylthiazol-2-yl)-2,5-diphenyltetrazolium bromide (MTT) assay to evaluate cell viability. A549 cells were seeded at a density of 10^5^ cells per well in a 96-well plate. The culture medium was replaced with a new medium containing 0, 1, 5, or 10 parts per billion (ppb) of PFAS. After incubating the cells with or without PFAS for either 12 or 24 hours, an MTT assay was conducted to evaluate the cytotoxic effects of PFAS on A549 cells. For the MTT assay, 5 mg/mL MTT (3-(4,5-Dimethylthiazol-2-yl)-2,5-Diphenyltetrazolium Bromide) (Thermo Fisher Scientific, Waltham, MA, USA) was added to the culture medium at 10% of the medium to reach a final concentration of 0.5 mg/mL. The medium and MTT mixture was removed after 2 hours of incubation, and the resulting formazan crystals were dissolved in DMSO. Absorbance was measured at 570 nm wavelength using a FLUOStar Omega plate reader (BMG LabTech, Ortenberg, Germany) to quantify cell viability. Relative cell viability was calculated using the following formula:

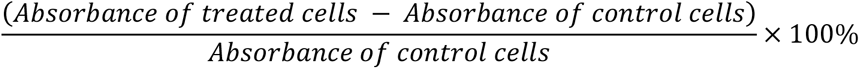

### 2.4 RNA Extraction and Quantitative Real-Time PCR

Total RNA was extracted from exposed and control cells using the RNeasy Plus Kit (QIAGEN, Germantown, MD, USA) according to the manufacturer’s instructions. Briefly, cells were lysed and homogenized in the supplied lysis buffer, and genomic DNA was removed using the integrated gDNA Eliminator spin column. The resulting lysate was then applied to an RNeasy spin column, washed, and eluted with RNase-free water to obtain purified total RNA. The extracted total RNA was reverse transcribed into cDNA using the RevertAid RT Reverse Transcription Kit (Thermo Fisher Scientific, Waltham, MA, USA) according to the manufacturer’s protocol. All primers used for quantitative real-time PCR were purchased from Integrated DNA Technologies (IDT, Coralville, IA, USA). The cDNA samples were subsequently mixed with qPCR Master Mix and gene-specific primers, and quantitative real-time PCR was performed using the QuantStudio 3 Real-Time PCR System. Relative gene expression levels were calculated after normalization to the GAPDH gene, and all procedures were performed according to the manufacturers’ instructions.

### 2.5 Data analysis

Raw GC–MS data files were processed using Compound Discoverer™ version 3.3.3.2 (Thermo Fisher Scientific, Waltham, MA, USA) for peak extraction, peak alignment, deconvolution, compound annotation, and relative quantification. Compound identification was performed by matching mass spectra to the NIST Mass Spectral Library and was further supported by comparing experimentally determined linear retention indices (LRIs) with those calculated from a homologous series of n-alkane standards (C8–C20; Sigma-Aldrich, St. Louis, MO, USA) analyzed under identical chromatographic conditions. Prior to statistical analysis, peak areas were total ion current (TIC)-normalized and log_10_-transformed. Differential metabolite analysis was performed using Student’s t-test for pairwise comparisons, with statistical significance defined as p < 0.05 and a |log_2_ fold change| ≥ 1. To control for multiple comparisons, Benjamini–Hochberg false discovery rate (FDR) correction was applied where appropriate. Principal component analysis (PCA), volcano plots, heatmaps, and metabolite set enrichment analysis (MSEA) were performed using Compound Discoverer and MetaboAnalyst 6.0 to visualize metabolic variation and identify significantly altered volatile organic compounds (VOCs). For RT-qPCR analysis, cycle threshold (Ct) values were obtained using the QuantStudio™ 3 Real-Time PCR System (Applied Biosystems, Thermo Fisher Scientific, Waltham, MA, USA). Target gene expression was normalized to GAPDH, and relative expression levels were calculated using the 2^−ΔΔCt method, with untreated control samples serving as the calibrator. Data are presented as mean ± standard error of the mean (SEM). Statistical significance was evaluated using one-way analysis of variance (ANOVA) followed by Tukey’s multiple comparisons test, with p < 0.05 considered statistically significant.

## 3. Results

### 3.1 PFAS exposure alters VOC profiles

Principal component analysis (PCA) was used to assess global changes in VOC profiles following PFAS exposure (Fig. 1). In A549 cells, the first two principal components accounted for 33.8% (PC1) and 14.3% (PC2) of the total variance. The score plot showed distinct clustering of samples according to exposure time, with 12, 24, and 48 h samples forming separate groups (along PC1). Within each exposure time, PFAS-treated samples were clearly separated from the controls and exhibited concentration-dependent clustering across the 1, 5, and 10 ppb treatment groups (along PC2). The greatest separation from the control was observed at 10 ppb (Fig.1a). Similarly, the PCA score plot of BEAS-2B cells (Fig. 1b) showed clear separation of samples by exposure time, with PC1 and PC2 accounting for 48.9% and 13.7% of the total variance, respectively. Within each exposure time, PFAS-treated samples were separated from the controls, although clustering was tighter than that observed in A549 cells of treatment groups (Fig. 1b). Biological replicates clustered closely within each treatment group, indicating high analytical reproducibility.

**Fig. 1.**
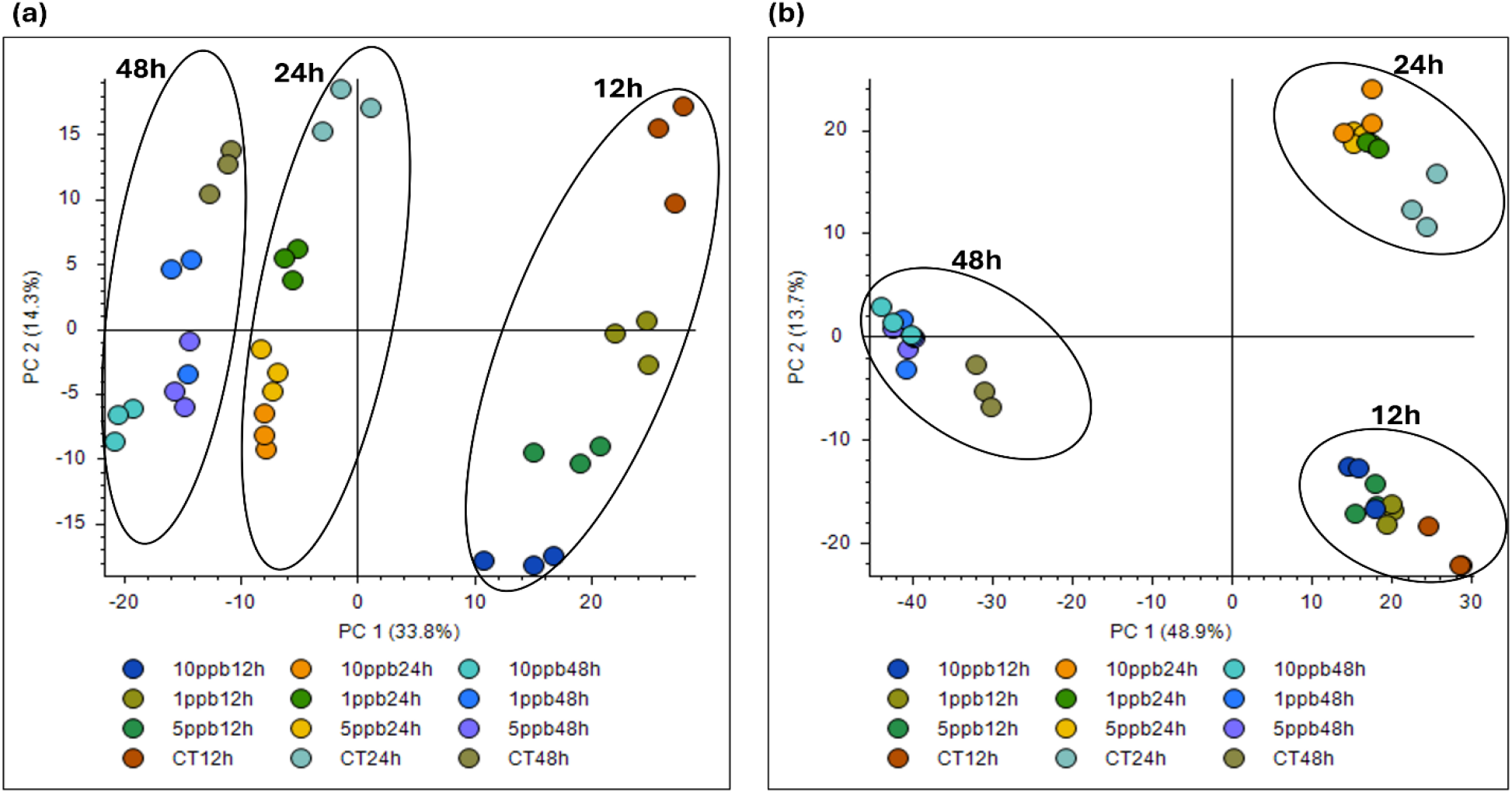
Principal Component Analysis (PCA) score plots of VOC profiles showing separation of PFAS-treated samples and controls (CT) for (a) A549 and (b) BEAS-2B cells across 12 h, 24, and 48 h and concentrations of 1ppb, 5ppb, and 10ppb. Each color set indicates biological replicates.

### 3.2 PFAS exposure alters VOC abundance and response patterns

Concentration-and time-dependent changes in VOC features were further supported by volcano plot analysis (Fig. 2a-f). Differentially abundant VOC features were identified using a threshold of p < 0.05 and |log_2_ fold change| ≥ 1. Significantly increased, decreased, and unchanged VOC features are shown in orange, green, and gray, respectively. In A549 cells, significantly altered VOC features were detected at all PFAS concentrations, with the greatest number observed at 10 ppb (Fig. 2a-c). A similar concentration-dependent trend was observed in BEAS-2B cells (Fig. 2d-f), although more significantly altered features were observed than in A549 cells. Time-dependent volcano plot analysis at 10 ppb PFAS revealed distinct temporal patterns in VOC abundance (Fig. 3a-f). In A549 cells, significantly upregulated VOC features increased over time and were most abundant at 48 h, whereas significantly downregulated VOC features decreased from 12 to 48 h. In BEAS-2B cells, the greatest number of significantly upregulated VOC features was observed at 12 h, while significantly downregulated VOC features were most abundant at the 48 h time point.

**Fig. 2.**
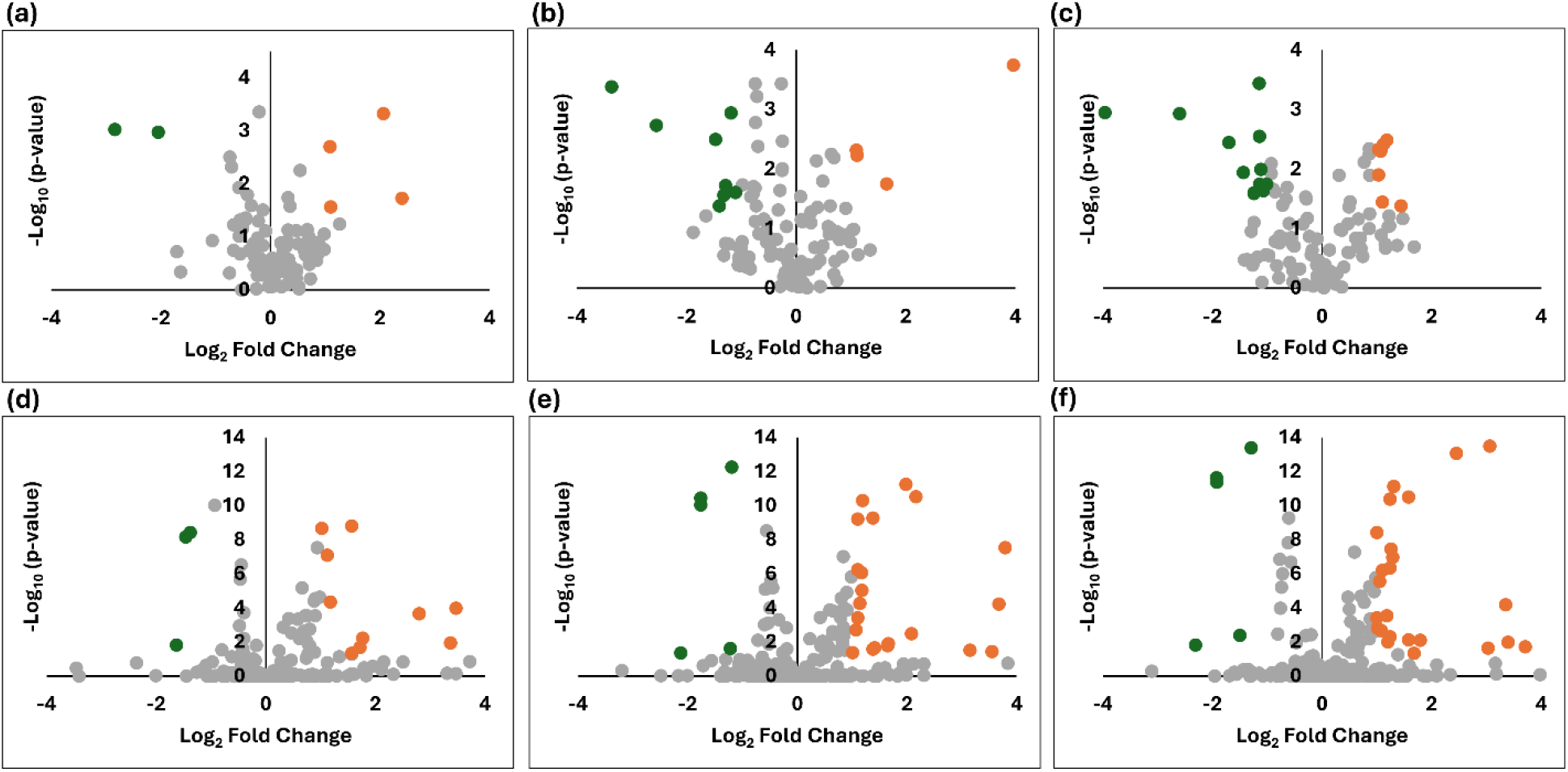
Concentration-dependent volcano plots for VOC features following 12 h PFAS exposure to 1 ppb, 5 ppb, and 10 ppb in A549 cells (a-c) and BEAS-2B cells (d-f). Green dots indicate significantly downregulated features, orange points indicate significantly upregulated VOC features, and gray points represent non-significant features. Statistical significance was defined as p-value < 0.05 and |log_2_ fold change| ≥ 1. Volcano plots for the 24 h and 48 h PFAS exposures are provided in Fig. S1 and Fig. S2, respectively.

**Fig. 3.**
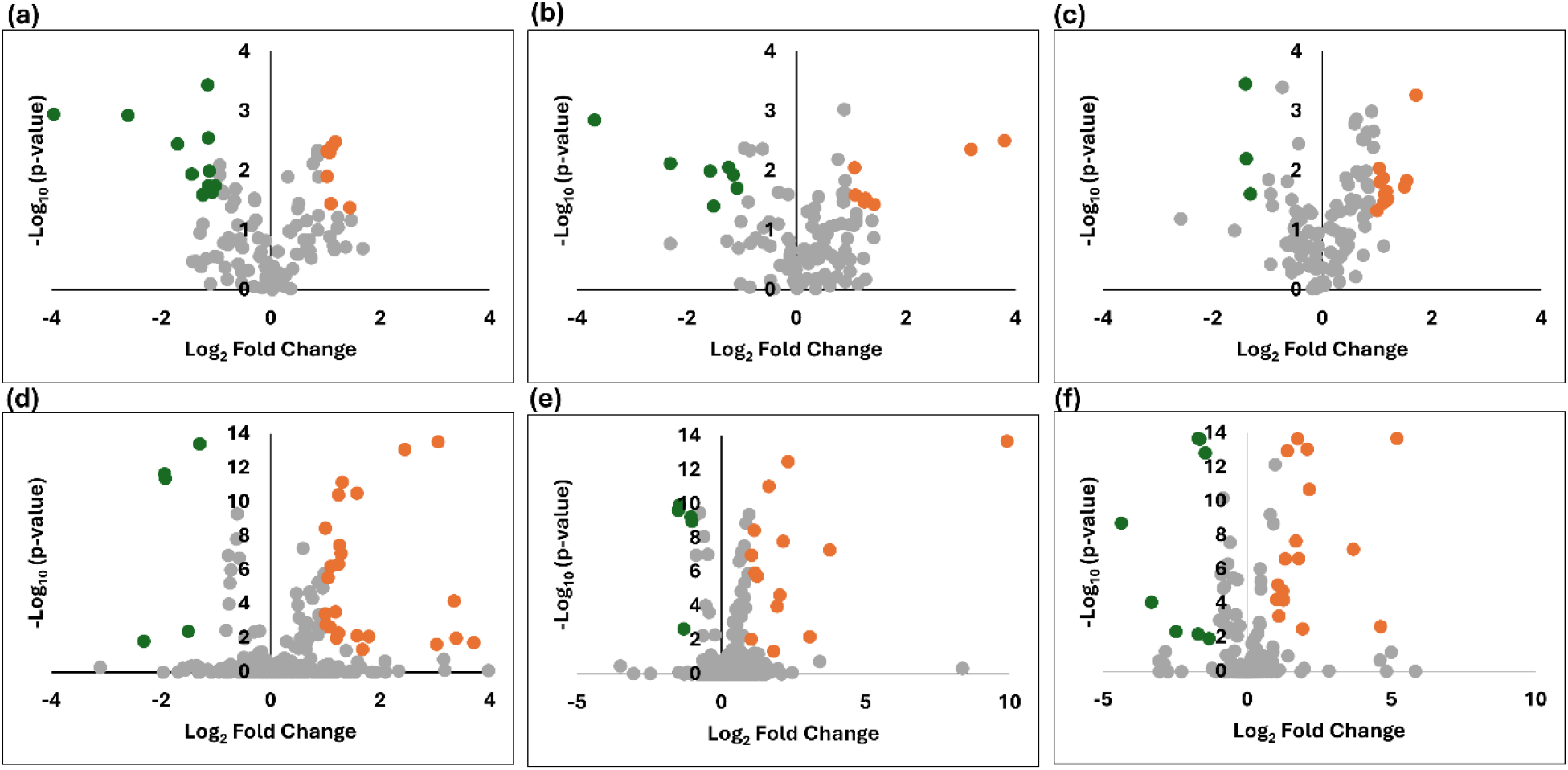
Time-dependent volcano plots of VOC features following 10 ppb PFAS exposure at 12 h, 24 h, and 48 h in A549 cells (a-c) and BEAS-2B cells (d-f). Green points represent significantly decreased VOC features, orange points represent significantly increased VOC features, and gray points represent non-significant features. Statistical significance was defined as p-value < 0.05 and |log_2_ fold change| ≥ 1. Volcano plots for the 1 ppb and 5 ppb PFAS exposures are provided in Fig. S3 and Fig. S4, respectively.

Hierarchical clustering of the putatively identified VOCs further revealed distinct response patterns across PFAS concentrations and exposure times in both A549 and BEAS-2B cells (Fig. 4). In A549 cells, several VOCs, including acetoin, (E)-2-nonenal, acetone, 2-nonanone, (E)-2-octenal, and the 2,4-alkadienals, generally exhibited higher relative abundances with increasing PFAS concentration and at later exposure time. In contrast, compounds such as butanal, pentanal, 2-butanone, and 1-hexanol exhibited relatively higher abundances in the control and early-exposure groups and lower relative abundances after PFAS treatment. A similar overall pattern was observed in BEAS-2B cells, where acetone, (E)-2-nonenal, (E)-2-decenal, 5-nonanone, and (E)-2-octenal exhibited progressively higher relative abundances, particularly at higher PFAS concentrations and at 48 h. Conversely, pentanal, acetoin, 1-hexanol, butanal, and 2-butanone showed comparatively higher abundances in the control and early-exposure groups, with their relative abundances decreasing following PFAS exposure. Overall, the heatmaps demonstrate that PFAS exposure was associated with distinct temporal- and concentration-dependent alterations in the profiles of putatively identified VOCs, although the response patterns differed between A549 and BEAS-2B cells. The identities, library matching scores, and corresponding statistical p-values for identified VOCs under each condition are provided in Table S1 and Table S2.

**Fig. 4.**
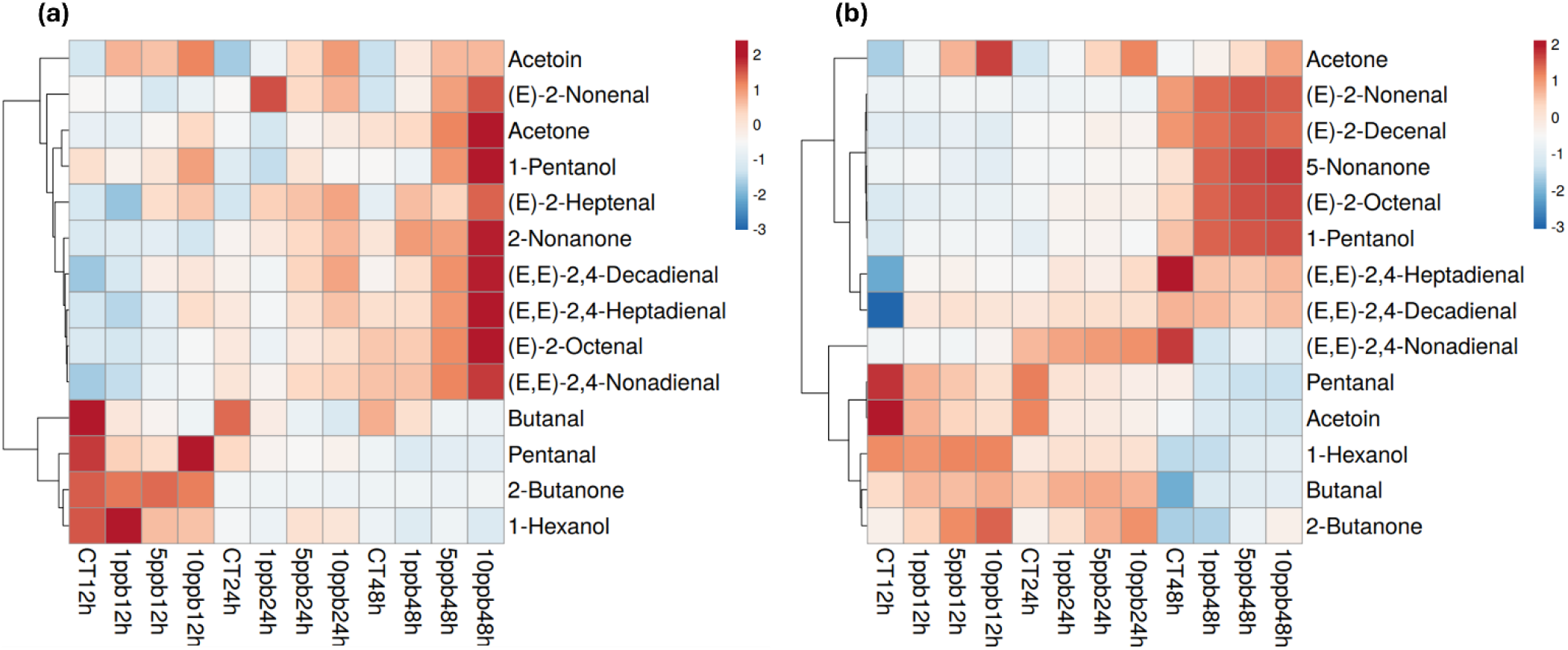
Hierarchical clustering heatmaps of differentially expressed volatile organic compounds (VOCs) in (a) A549 and (b) BEAS-2B cells following PFAS exposure. Rows represent individual putatively identified VOCs, and columns represent the mean VOC abundance for each treatment group across exposure times and PFAS concentrations. Data were log-transformed, and heatmaps were generated using the ClustVis web service with hierarchical clustering based on correlation distance and average linkage. The color scale represents relative abundance: red indicates increased abundance, blue indicates decreased abundance, and white indicates no significant change relative to the mean normalized abundance.

### 3.3 Metabolite set enrichment analysis

To evaluate the biological meaning of the significantly altered VOCs following PFAS exposure, metabolite set enrichment analysis was performed using MetaboAnalyst 6.0 over-representation analysis (ORA) against the RaMP-DB metabolite set database. The enrichment profiles were highly similar between A549 and BEAS-2B cells, with the same metabolite sets consistently identified in both cell lines (Fig. 5a and Fig. 5b). Among the enriched metabolite sets, ketone body metabolism, ketogenesis and ketolysis exhibited relatively higher enrichment ratios. In contrast, cytochrome P450-mediated Phase I functionalization of compounds and biological oxidations showed comparatively lower enrichment ratios.

**Fig. 5.**
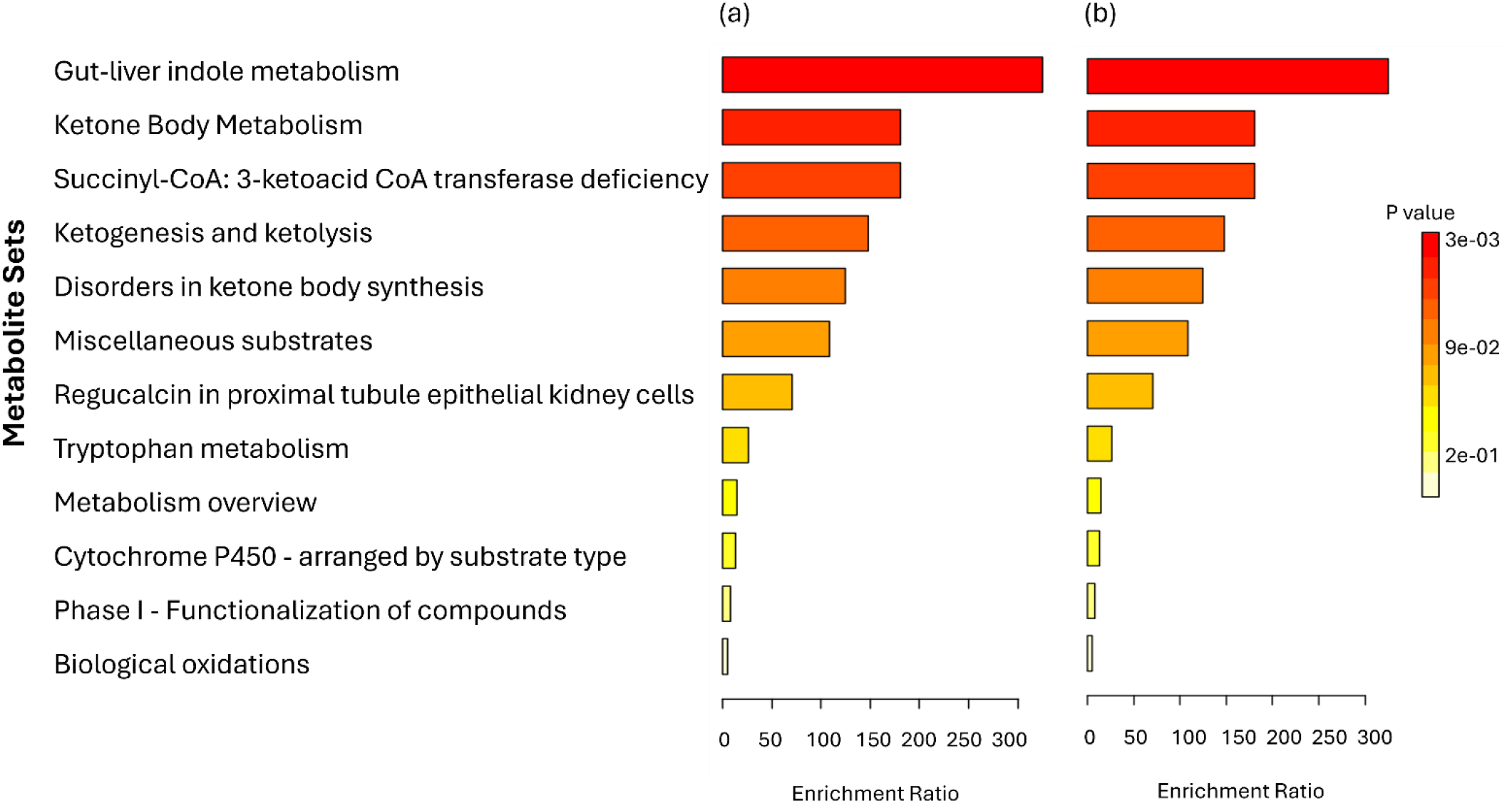
Metabolite set enrichment analysis following PFAS exposure in (a) A549 and (b) BEAS-2B cells. Bar length represents the enrichment ratio, while bar color indicates statistical significance (p-value), with darker red indicating lower p-values.

### 3.4 Effects of PFAS pulmonary exposure on xenobiotic metabolism and aldehyde detoxification gene expressions

To further investigate the molecular responses associated with the altered VOC profiles following PFAS exposure, we evaluated the expression of selected genes involved in xenobiotic metabolism and aldehyde detoxification. The selection of cytochrome P450 (CYP) and aldehyde dehydrogenase (ALDH) genes was guided by the observed VOC profiles, which included multiple aldehydes and other oxidation-related VOCs. Accordingly, the relative mRNA expression of CYP1A1, CYP2E1, ALDH1A1, and ALDH2 was assessed in A549 and BEAS-2B cells following 24 h of exposure (Fig. 6). In A549 cells, ALDH1A1 mRNA expression was significantly decreased (0.7-fold), with no significant decrease in ALDH2 expression (0.9-fold). CYP1A1 expression increased (1.4-fold), whereas CYP2E1 decreased (0.3-fold) relative to the control (1.0), with no significant difference. In BEAS-2B cells, CYP1A1 expression was significantly decreased (0.9-fold), while ALDH1A1, ALDH2, and CYP2E1 expression decreased, but the differences were not statistically significant.

**Fig. 6.**
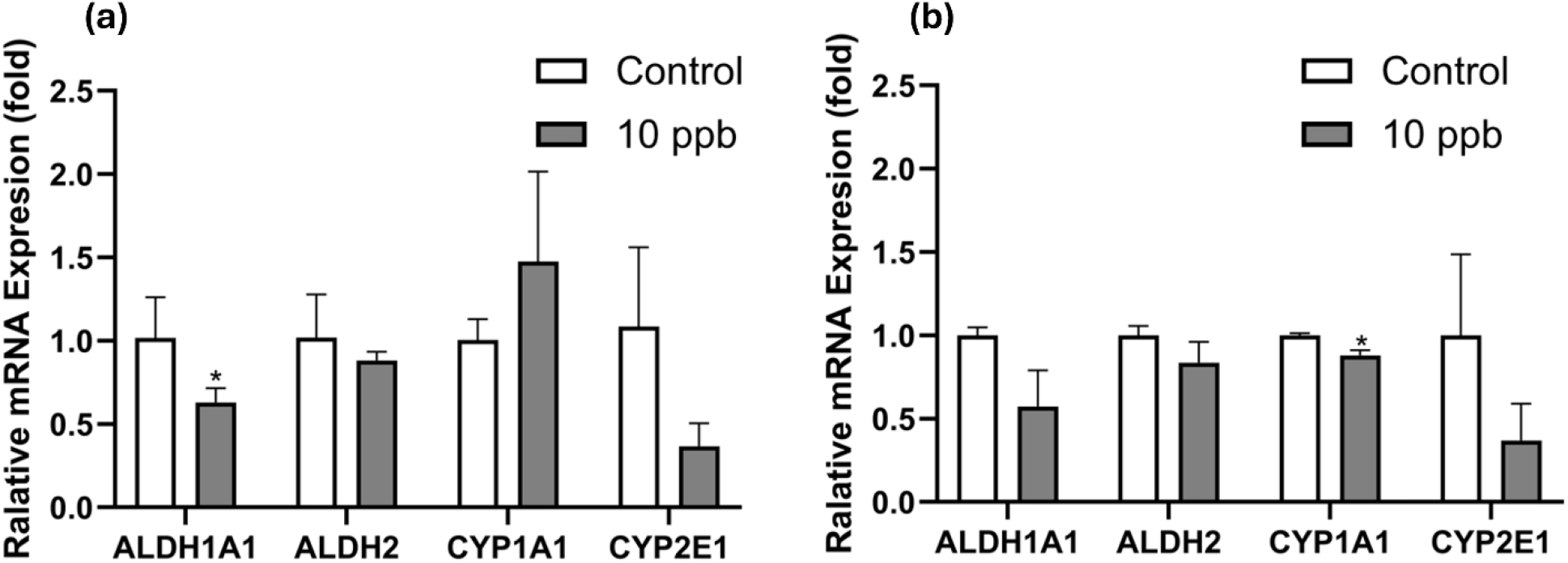
Relative mRNA expressions of ALDH1A1, ALDH2, CYP1A1, and CYP2E1 in (a) A549 and (b) BEAS-2B cells following 24 h PFAS exposure. Gene expression levels were normalized to the control group (GAPDH) and are presented as mean ± SD (n = 3). Statistical significance was determined using one-way ANOVA followed by Tukey’s post hoc test (\**p* < 0.05).

## 4. Discussion

The present study demonstrated that exposure of human lung epithelial cells to environmentally relevant PFAS concentrations and sub-lethal concentrations altered volatile organic compound (VOC) profiles in a concentration- and time-dependent manner. Given the current lack of defined PFAS-specific VOC biomarkers in lung epithelial cells,^46,47^ we integrated untargeted analysis using HS-SPME GC-Orbitrap MS with multivariate analysis, metabolite set enrichment, and qPCR to delineate metabolic responses across PFAS concentrations (1, 5, and 10 ppb) at exposure times (12, 14, and 48 h) in both cell types.

Consistent with prior toxicometabolomic studies in other cell models,^48-52^ our findings confirm that PFAS exposure induces significant and non-monotonic metabolic changes. PCA showed the distinct separation by exposure time, suggesting that exposure duration was a major determinant of the metabolic response. Within each time point, controls were separated from PFAS-exposed samples. At the same time, volcano plots revealed that the number of significantly altered VOC features varied across exposures at 1, 5, and 10 ppb across time points. At 1 ppb, fewer significantly dysregulated VOC features were observed in both cell models, suggesting that even low PFAS levels may trigger an early, limited metabolic response due to the activation of specific pathways. Prior work supports this, showing that low-dose exposures to persistent organic pollutants can elicit non-monotonic or subtle toxicological effects.^53^ The upregulation and downregulation of several statistically significant VOC features after a longer duration of exposure (48 h) at a higher concentration (10 ppb) suggest late-phase responses consistent with enhanced cellular stress and broader activation or inactivation of metabolic pathways leading to a more diverse and detectable array of metabolites.^54^ This pattern suggests that PFAS exposure produces dynamic metabolic remodeling rather than a simple linear dose-response. Such non-uniform responses are biologically plausible as PFAS can simultaneously activate adaptive stress responses and disrupt metabolic pathways, with the relative contribution of each process changing over time.^55^

The altered VOC patterns were dominated by chemically and biologically meaningful classes, including aldehydes, ketones, and alcohols. These compound classes provide insight into the metabolic processes most affected by PFAS exposure and suggest involvement of oxidative stress, lipid peroxidation, ketone body metabolism, and xenobiotic-related metabolism^56^. The increased coordinated changes among several VOCs, including acetone, 2-nonanone, 5-nonanone, (E)-2-octenal, (E)-2-nonenal, (E)-2-decenal, and related 2,4-alkadienals such as 2,4-heptadienal, 2,4-nonadienal, and 2,4-decadienal in both cells, represent two major biochemical responses: altered energy metabolism and oxidative damage to membrane lipids. The ketone-containing VOCs such as acetone, 2-nonanone, and 5-nonanone are associated with altered fatty acid β-oxidation, ketone body metabolism, and changes in cellular energy metabolism. Their abundance suggests that PFAS exposure shifts cellular metabolism toward greater lipid utilization, consistent with the enrichment of ketone body metabolism, ketogenesis, and ketolysis observed in the pathway analysis. Similar metabolic disturbances have been reported in PFAS studies that demonstrate that PFAS exposure disrupts cellular energy homeostasis and lipid metabolism.^57-60^ Also, the increased abundance of (E)-2-octenal, (E)-2-nonenal, (E)-2-decenal, and related 2,4-alkadienals is a well-established product of lipid peroxidation generated during reactive oxygen species (ROS)-mediated oxidation of membrane polyunsaturated fatty acids, although the present study does not directly quantify ROS or lipid peroxidation products. Lipid peroxidation occurs when oxidants attack membrane lipids, producing reactive aldehydes that serve as indicators of oxidative stress and membrane damage.^61^ This is further supported by previous studies showing that PFAS exposure promotes oxidative stress and membrane dysfunction^62,63^, supporting the biological plausibility of the aldehyde alterations observed in the present study. In contrast, several VOCs, including acetoin, butanal, pentanal, 2-butanone, and 1-hexanol, exhibited reduced abundance following PFAS exposure. This suggests that PFAS exposure disrupts interconnected metabolic networks rather than isolated pathways, leading to different chemical classes of VOCs.

The supplementary VOC identification tables (Tables S1 and S2) further support the compound-level alterations described above. Most putatively identified VOCs were common to both A549 and BEAS-2B cells, with only minor differences in the specific compounds detected. However, BEAS-2B exhibited statistically significant changes for many of the same VOCs across a greater number of PFAS concentrations and exposure times than A549 cells. These observations complement the heatmap analyses by demonstrating that PFAS exposure elicited comparable VOC profiles in both cell lines while differing in the statistical significance of the responses across exposure conditions.

Pathway enrichment analysis further supported the metabolic alterations observed in the VOC profiles. Enrichment of ketone body metabolism, ketogenesis, and ketolysis is consistent with the changes in ketone-containing VOCs, whereas enrichment of biological oxidation supports the accumulation of lipid peroxidation-derived aldehydes. The enrichment of these metabolite sets suggests that PFAS exposure affects multiple interconnected metabolic processes rather than isolated biochemical reactions. The enrichment of biological oxidation, cytochrome P450 metabolism, and Phase I functionalization pathways further supports the hypothesis that PFAS exposure activates cellular detoxification mechanisms. Phase I metabolism is primarily mediated by cytochrome P450 enzymes, which catalyze oxidation, hydroxylation, and other reactions that increase the polarity of endogenous and xenobiotic compounds. Although PFAS are highly resistant to direct enzymatic metabolism due to the strength of the carbon-fluorine bond, PFAS exposure can alter the expression and activity of xenobiotic-metabolizing enzymes, thereby influencing the metabolism of endogenous substrates and other environmental chemicals. Consequently, the enrichment of these metabolite sets likely reflects secondary metabolic responses to PFAS exposure rather than direct PFAS biotransformation.

The altered VOC profiles observed following PFAS exposure, particularly the presence of multiple oxidation-related VOCs, motivated the further investigation of the molecular effects of PFAS exposure. We evaluated the expression of CYP1A1 and CYP2E1, two phase I cytochrome P450 enzymes involved in the metabolism of xenobiotics (environmental contaminants).^64^ Following 24 h of PFAS exposure, CYP1A1 expression was significantly decreased in BEAS-2B cells, whereas A549 cells exhibited a non-significant increase. The significant decrease in CYP1A1 expression observed in BEAS-2B cells is consistent with a previous study reporting reduced CYP1A1 expression in human bronchial epithelial cells following 24 h exposure to perfluorohexanoic acid (PFHxA),^65^ suggesting that PFAS can suppress CYP1A1 expression in normal airway epithelial cells. The non-significant increase in CYP1A1 observed in A549 cells suggests that PFAS-induced responses are cell-type dependent and may differ between transformed and non-transformed lung epithelial cells. CYP2E1 expression decreased in both cell types. Similarly, a transcriptomic analysis of 33 PFAS in HepaRG cells demonstrated that PFAS are not universal inducers of cytochrome P450 enzymes. Within these altered xenobiotic metabolic pathways, CYP1A1 and CYP2E1 were downregulated in HepaRG cells.^66^ Therefore, the reduced expression of CYP1A1 and CYP2E1 following PFAS exposure suggests diminished capacity for phase I xenobiotic metabolism in the airway, thereby reducing the cellular capacity for phase I detoxification of xenobiotics and other small organic molecules. Such alterations may increase reactive oxygen species production, promoting oxidative stress, inflammation, and epithelial barrier dysfunction, similar to patterns reported following particulate matter 2.5 μm (PM_2.5_) induced pulmonary injury.^67-69^

Furthermore, the detection of multiple aldehydes among the altered VOCs guided our evaluation of ALDH1A1 and ALDH2, two aldehyde dehydrogenase enzymes. ALDH enzymes play an essential role in cellular defense against oxidative damage by metabolizing reactive aldehydes generated during lipid peroxidation into less toxic carboxylic acids.^70^ Following PFAS exposure, ALDH1A1 expression was significantly reduced in A549 cells, whereas ALDH2 showed a modest decrease in both cell lines. Therefore, reduced ALDH1A1 expression, together with the modest decline in ALDH2, may promote the accumulation of reactive aldehydes, which are toxic byproducts of lipid peroxidation.^71,72^ Consequently, reduced ALDH1A1 expression may diminish the ability of airway epithelial cells to manage oxidative stress, a key mechanism underlying PFAS toxicity,^63,73^ thereby contributing to lung dysfunction and tissue damage. Hence, the reduced ALDH expression observed following PFAS exposure may also contribute to the altered aldehyde-containing VOC profiles detected in this study.

An important implication of the present findings is that VOC profiling detected biochemical alterations before overt cytotoxicity. Although traditional cytotoxicity assays indicated reduced cell viability at 12 h and 24 h post-exposure (Fig. S5), these changes were not statistically significant. Similar studies have demonstrated that metabolomic biomarkers can reveal pathological changes before they become detectable through conventional bioassays.^74-78^ This supports the use of volatilomics as a sensitive approach for identifying early cellular responses to chemical exposure. VOCs are downstream products of multiple biochemical processes. Therefore, changes in VOC profiles may occur before large changes in cell viability are observed.

In conclusion, this study demonstrates that environmentally relevant PFAS exposure induces concentration- and time-dependent alterations in the volatile metabolome of lung epithelial cells. High-resolution mass spectrometry identified changes in aldehydes, ketones, and alcohols, indicating perturbations in lipid metabolism, mitochondrial energy metabolism, and xenobiotic metabolism. Both cell lines exhibited early and more pronounced metabolic perturbations at higher concentrations. Pathway enrichment analysis further implicated ketone body metabolism, biological oxidation, and related metabolic processes. Complementary qPCR analysis showed altered expression of genes involved in xenobiotic metabolism (CYP1A1 and CYP2E1) and aldehyde detoxification (ALDH1A1 and ALDH2), supporting the metabolomic findings and suggesting impaired cellular detoxification and oxidative stress responses following PFAS exposure. Collectively, these findings demonstrate that integrating volatile metabolomics with pathway enrichment and targeted gene expression analysis provides a comprehensive approach for characterizing early metabolic responses to PFAS exposure. The identified VOC signatures represent potential biomarkers for PFAS exposure while providing new mechanistic insight into PFAS-induced metabolic perturbations in lung epithelial cells.

## Author Contributions

**Patrica-Ivy Agorsor**: Data curation, formal analysis, investigation, methodology, software, writing – original draft. **Ta-Chun Lin**: Formal analysis, investigation, methodology, writing – original draft. **Yue-Wern Huang**: Conceptualization, resources, software, supervision, validation, writing – review/editing. **Michael Eze**: Conceptualization, data curation, formal analysis, investigation, methodology, project administration, resources, software, supervision, validation, visualization, writing – review/editing.

## Funding

This study was supported in part by the Missouri S&T’s Spring 2025 Ignition Grant Initiative. The funders have no role in the study design, collection, analysis and interpretation of data, writing of the report, and decision to submit the article for publication.

## Notes

### Competing Interest Statement

The authors have declared no competing interest.

