## Supplementary Materials for "Volatile Metabolic Signatures Reveal Early Pulmonary Responses to PFAS Exposure"

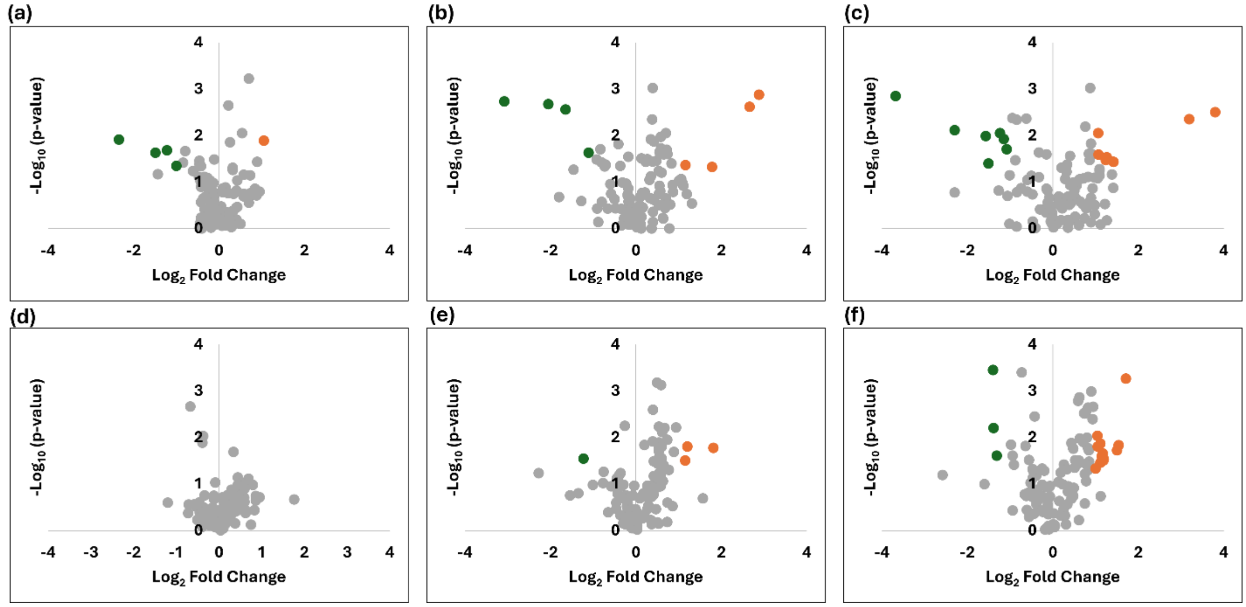


**Fig. S1.** Concentration-dependent volcano plots for VOC features following 24 h PFAS exposure to 1 ppb, 5 ppb, and 10 ppb (a-c) and 48 h (d-f) in A549 cells. Green dots indicate significantly downregulated features, orange points indicate significantly upregulated VOC features, and gray points represent non-significant features. Statistical significance was defined as p-value < 0.05 and |$\log_2$ fold change| ≥ 1.

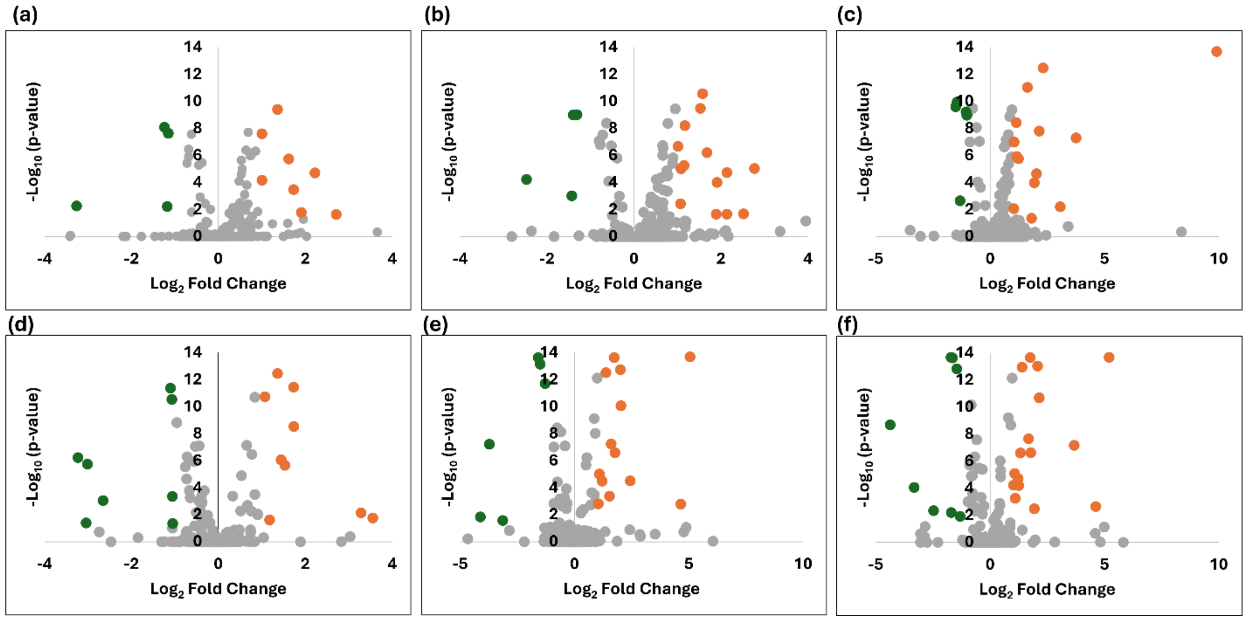


**Fig. S2.** Concentration-dependent volcano plots for VOC features following 24 h PFAS exposure to 1 ppb, 5 ppb, and 10 ppb (a-c) and 48 h (d-f) in BEAS-2B cells. Green dots indicate significantly downregulated features, orange points indicate significantly upregulated VOC features, and gray points represent non-significant features. Statistical significance was defined as p-value < 0.05 and |log$_2$ fold change| ≥ 1.

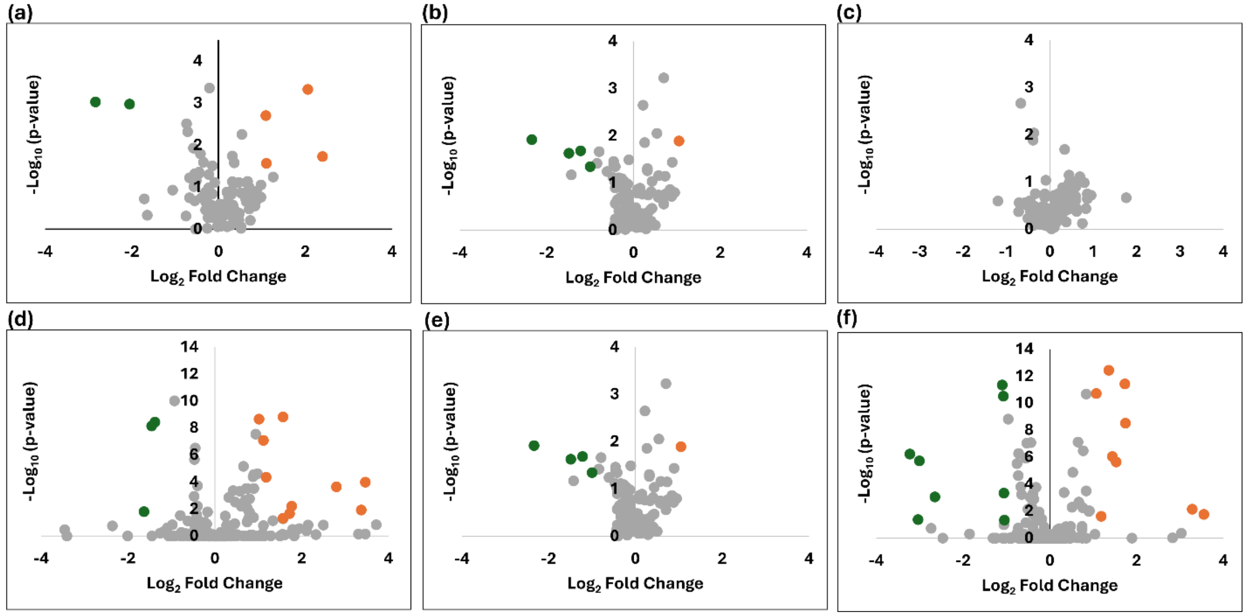


**Fig. S3.**Time-dependent volcano plots of VOC features following 1 ppb PFAS exposure at 12 h, 24 h, and 48 h in A549 cells (a-c) and BEAS- 2B cells (d-f). Green points represent significantly decreased VOC features, orange points represent significantly increased VOC features, and gray points represent non-significant features. Statistical significance was defined as p-value < 0.05 and |log2 fold change| ≥ 1.

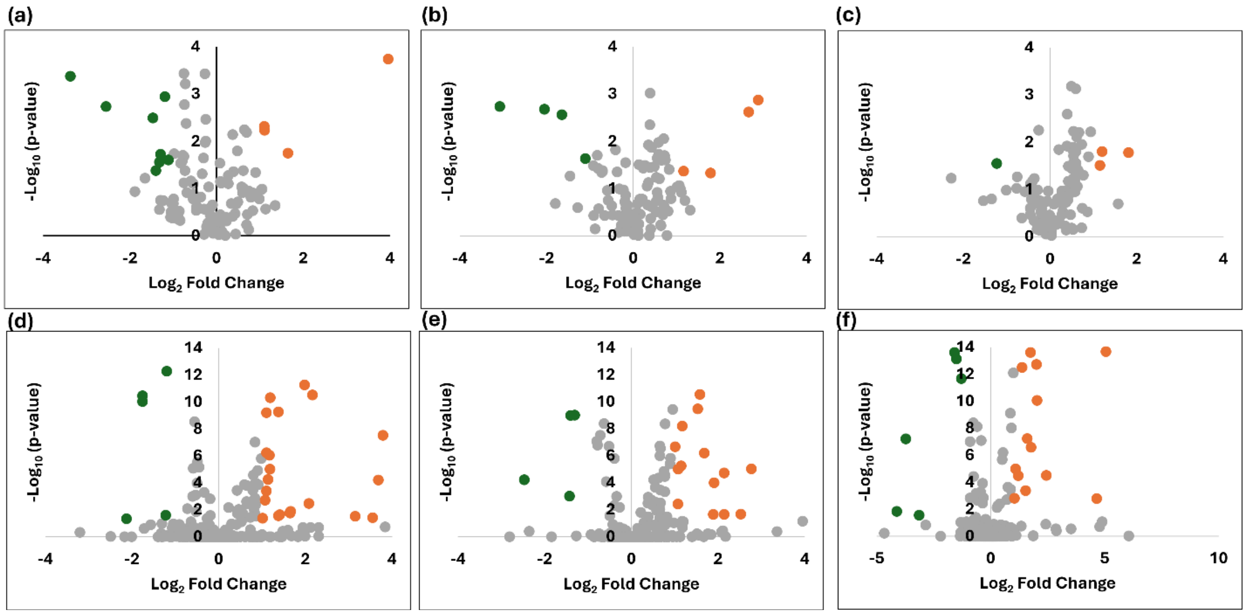


**Fig. S4.** Time-dependent volcano plots of VOC features following 5 ppb PFAS exposure at 12 h, 24 h, and 48 h in A549 cells (a-c) and BEAS- 2B cells (d-f). Green points represent significantly decreased VOC features, orange points represent significantly increased VOC features, and gray points represent non-significant features. Statistical significance was defined as p-value < 0.05 and $|\log_2 \text{ fold change}| \geq 1$.

**Table S1.** Key putatively identified volatile organic compounds following PFAS exposure in A549 cells. The table includes molecular weight, retention time (RT), NIST library match(total) score, and the corresponding p-values for each exposure condition. Statistical significance was determined using one-way ANOVA followed by Tukey's multiple comparison test.

| | | | | | | 12h | | | 24h | | | 48h | | |
|---|---|---|---|---|---|---|---|---|---|---|---|---|---|---|
| **#** | **Compound** | **Molecular Weight** | **RT (min)** | **Formula** | **Total Score** | **1 ppb** | **5 ppb** | **10 ppb** | **1 ppb** | **5 ppb** | **10 ppb** | **1 ppb** | **5 ppb** | **10 ppb** |
| 1 | Acetone | 58.041 | 3.433 | $C_3H_6O$ | 97.8 | - | - | - | - | - | - | - | 0.001 | 0.031 |
| 2 | Butanal | 72.057 | 5.942 | $C_4H_8O$ | 97.1 | 0.001 | 0.002 | 0.001 | 0.023 | 0.002 | 0.008 | - | - | - |
| 3 | 2-Butanone | 72.057 | 3.180 | $C_4H_8O$ | 100.0 | 0.012 | 0.003 | - | - | - | - | - | - | - |
| 4 | Pentanal | 86.073 | 6.959 | $C_5H_{10}O$ | 78.3 | 0.016 | 0.048 | - | - | - | - | - | - | - |
| 5 | Acetoin | 88.052 | 5.412 | $C_4H_8O_2$ | 80.0 | - | - | - | - | 0.040 | 0.029 | - | 0.020 | 0.009 |
| 6 | 1-Pentanol | 88.088 | 5.210 | $C_5H_{12}O$ | 79.6 | - | - | 0.013 | - | 0.001 | 0.050 | - | - | 0.040 |
| 7 | 1-Hexanol | 102.104 | 7.128 | $C_6H_{14}O$ | 70.9 | 0.014 | 0.016 | - | - | - | - | - | - | - |
| 8 | (E,E)-2,4-Heptadienal | 110.073 | 7.012 | $C_7H_{10}O$ | 76.1 | - | - | 0.036 | 0.040 | - | - | - | 0.027 | 0.025 |
| 9 | (E)-2-Octenal | 126.104 | 7.844 | $C_8H_{14}O$ | 79.3 | - | - | - | - | - | - | - | 0.003 | 0.035 |
| 10 | (E,E)-2,4-Nonadienal | 138.104 | 8.038 | $C_9H_{14}O$ | 78.5 | - | 0.042 | 0.004 | - | - | - | - | 0.001 | 0.029 |
| 11 | (E)-2-Nonenal | 140.120 | 8.082 | $C_9H_{16}O$ | 76.5 | - | - | - | 0.014 | 0.031 | - | - | 0.012 | 0.017 |
| 12 | (E)-2-Decenal | 154.135 | 8.682 | $C_{10}H_{18}O$ | 78.7 | - | - | - | - | - | - | 0.040 | 0.031 | 0.019 |
| 13 | 2-Nonanone | 142.135 | 8.628 | $C_9H_{18}O$ | 98.9 | - | - | - | - | 0.032 | 0.100 | - | 0.032 | 0.033 |
| 14 | (E,E)-2,4-Decadienal | 152.120 | 9.271 | $C_{10}H_{16}O$ | 70.9 | - | - | 0.013 | - | 0.020 | 0.033 | - | 0.006 | 0.019 |

(-): Not statistically significant

**Table S2.** Key putatively identified volatile organic compounds following PFAS exposure in BEAS-2B cells. The table includes molecular weight, retention time (RT), NIST library match(total) score, and the corresponding p-values for each exposure condition. Statistical significance was determined using one-way ANOVA followed by Tukey's multiple comparison test.

| | | | | | | 12h | | | 24h | | | 48h | | |
|---|---|---|---|---|---|---|---|---|---|---|---|---|---|---|
| **#** | **Compound** | **Molecular Weight** | **RT (min)** | **Formula** | **Total Score** | **1 ppb** | **5 ppb** | **10 ppb** | **1 ppb** | **5 ppb** | **10 ppb** | **1 ppb** | **5 ppb** | **10 ppb** |
| 1 | Acetone | 58.041 | 3.433 | $C_3H_6O$ | 96.0 | 0.001 | 0.001 | 0.001 | 0.001 | 0.001 | 0.001 | - | 0.002 | 0.001 |
| 2 | Butanal | 72.057 | 5.942 | $C_4H_8O$ | 75.3 | 0.002 | 0.002 | 0.001 | 0.011 | 0.003 | 0.050 | 0.001 | 0.001 | 0.001 |
| 3 | 2-Butanone | 72.057 | 3.180 | $C_4H_8O$ | 98.8 | 0.006 | 0.001 | 0.001 | - | 0.001 | 0.001 | - | 0.001 | 0.001 |
| 4 | Pentanal | 86.073 | 6.959 | $C_5H_{10}O$ | 90.3 | - | - | - | - | - | - | - | 0.040 | 0.021 |
| 5 | Acetoin | 88.052 | 5.412 | $C_4H_8O_2$ | 80.0 | 0.001 | 0.001 | 0.001 | 0.001 | 0.001 | 0.001 | 0.007 | 0.001 | 0.003 |
| 6 | 1-Pentanol | 88.088 | 5.210 | $C_5H_{12}O$ | 72.0 | 0.017 | 0.009 | 0.007 | 0.004 | 0.001 | 0.019 | 0.001 | 0.001 | 0.001 |
| 7 | 1-Hexanol | 102.104 | 7.128 | $C_6H_{14}O$ | 79.7 | - | - | - | - | - | - | - | 0.018 | 0.003 |
| 8 | (E,E)-2,4-Heptadienal | 110.073 | 7.012 | $C_7H_{10}O$ | 72.7 | 0.001 | 0.001 | 0.001 | - | - | - | 0.042 | 0.028 | - |
| 9 | (E)-2-Octenal | 126.104 | 7.844 | $C_8H_{14}O$ | 93.4 | - | 0.018 | 0.010 | 0.031 | 0.003 | 0.015 | 0.001 | 0.001 | 0.001 |
| 10 | (E,E)-2,4-Nonadienal | 138.104 | 8.038 | $C_9H_{14}O$ | 71.7 | - | - | - | - | - | - | 0.001 | 0.002 | 0.001 |
| 11 | (E)-2-Nonenal | 140.120 | 8.082 | $C_9H_{16}O$ | 78.1 | - | - | - | - | - | - | 0.001 | 0.001 | 0.001 |
| 12 | (E)-2-Decenal | 154.135 | 8.682 | $C_{10}H_{18}O$ | 78.7 | - | - | - | - | - | - | 0.040 | 0.031 | 0.019 |
| 13 | 5-Nonanone | 142.135 | 8.641 | $C_9H_{18}O$ | 75.3 | - | - | - | - | - | - | 0.010 | 0.002 | 0.001 |
| 14 | (E,E)-2,4-Decadienal | 152.120 | 9.271 | $C_{10}H_{16}O$ | 74.6 | 0.073 | 0.022 | 0.050 | - | - | - | - | - | - |

(-): Not statistically significant

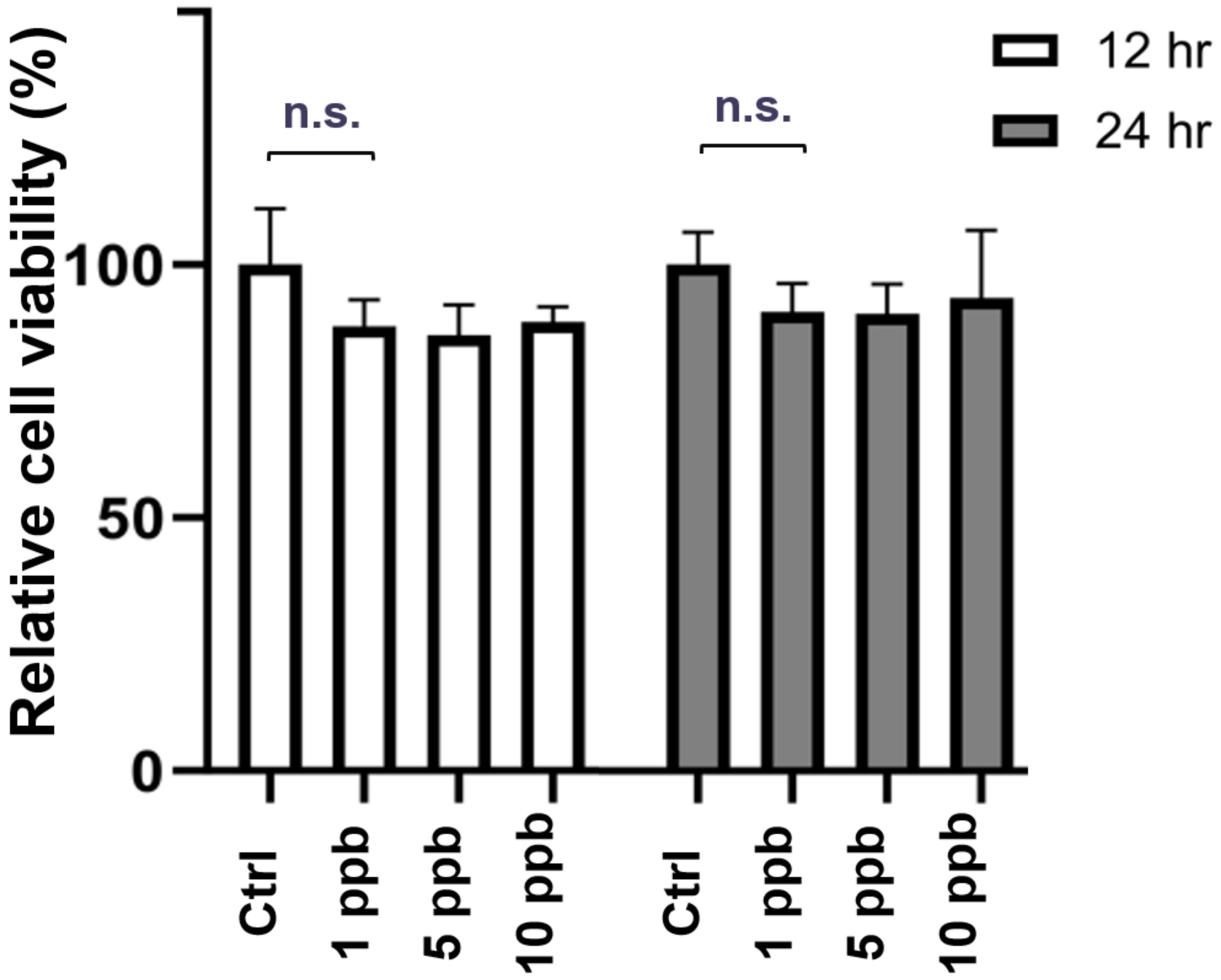


**Fig. S5.** Percentage relative viability at 12 h and 24 h of human alveolar basal epithelial cells (A549) exposed to varying concentrations of PFAS.
